# Self-perceived middle-distance race pace is faster in Advanced Footwear Technology spikes

**DOI:** 10.1101/2023.10.25.564056

**Authors:** Montgomery Bertschy, Victor Rodrigo-Carranza, Ethan W.C. Wilkie, Laura A. Healey, Jeremy Noble, Wayne J. Albert, Wouter Hoogkamer

**Affiliations:** Department of Kinesiology, University of Massachusetts, Amherst, MA, USA; Sports Performance Research Group (GIRD) University of Castilla-La Mancha, Toledo, Spain; University of New Brunswick, Fredericton, NB, Canada; PUMA SE, Innovation, Somerville, MA, USA

**Author notes:** These authors contributed equally to this work (Montgomery Bertschy, Victor Rodrigo-Carranza, Ethan W.C. Wilkie). **Corresponding Author**: Wouter Hoogkamer Integrative Locomotion Laboratory Department of Kinesiology University of Massachusetts, Amherst Amherst, MA, 01003-9258 USA.

**Keywords:** super spikes, track and field, running performance, carbon fiber plate, athletics

## Abstract

**Background:** Quantifying the potential benefits of advanced footwear technology (AFT) track shoes (i.e., “spikes”) in middle-distance events is challenging, because repeated maximal effort trials (as in sprinting) or aerobic running economy trials (as in long-distance running) are not feasible.

**Methods:** We introduce a novel approach to assess the benefits of AFT spikes, consisting of a series of 200 m runs at self-perceived middle-distance race pace with 10 min recovery and conducted four experiments to evaluate its validity, sensitivity, reproducibility, and utility.

**Results:** In experiment 1, participants ran 1.2% slower in spikes with 200 g added mass vs. control spikes, exactly equal to the known effects of shoe mass on running performance. In experiment 2, participants ran significantly faster in AFT prototype spikes vs. traditional spikes. In experiment 3, we compared two other AFT prototype spikes against traditional spikes, on three separate days. Group-level results were consistent across days, but our data indicates that at least two separate sessions are needed to evaluate individual responses. In experiment 4, participants ran significantly faster in two AFT spike models vs. traditional spikes (2.1% and 1.6%). Speed was similar between a third AFT spike model and the traditional spikes. These speed results were mirrored by changes in step length, as participants took significantly longer steps in the two faster AFT spike models (2.3% and 1.9%), while step length was similar between the other spikes.

**Conclusion:** Our novel, interval-based approach is a valid and reliable method to quantify differences between spikes at middle-distance running intensity.

## 1 Introduction

Once thought to be an unbreakable barrier on the track ^1^, the magical sub-4-minute mile mark stood strong for decades, until Sir Roger Bannister in 1954 completed the unthinkable and ran the 1609.344 meters in 3:59.4 seconds. In February 2023, 52 male professional, club, and collegiate level athletes broke the 4-minute mile barrier all within the span of an hour at Boston University’s indoor track facility. A similar phenomenon was observed in the women’s section, as three athletes broke the 4:30 mile barrier, and 17 ran sub-4:36, a mark some consider to be more equivalent to the men’s barrier.^2^ Over the past decades, the number of athletes breaking these barriers had been increasing gradually, but there has been a substantial increase since 2021. While the Boston University track has been known to be fast, another important common feature among these performances was the use of advanced footwear technology (AFT) track shoes (i.e., “spikes”), colloquially known as “super spikes”. Similar to the now common AFT marathon racing shoes (e.g., Nike Vaporfly), AFT track spikes feature a lightweight, compliant, and resilient midsole foam, often combined with a stiff midsole plate. While we have argued before that anecdotal performances should not be used as evidence on the efficacy of specific shoes ^3^, the widespread improvement across middle-distance running performances warrants a systematic, scientific controlled investigation.

The recent improvements on the track are similar to improvements in road running performances since 2017 which have been partly attributed to advances in road running shoes.^4–6^ While the adoption of AFT road shoes happened in parallel with the publication of a randomized crossover study showing 4% improvements in running economy,^7^ scientific evidence on potential benefits of AFT track spikes is currently limited to race results analyses (preprint: ^8^) and preliminary laboratory data.^9^ In an observational study, Willwacher et al.^8^ found that AFT track spikes had a greater effect on performance in middle-distance events (0.31 - 1.33%) than in sprint events (0.22 - 0.66%) for both men and women.

Traditionally, the effects of footwear on middle-distance running performance have been studied in the lab by measuring running economy, or on the track with time trials. Barnes and Kilding^10^ observed that running economy in traditional lightweight track spikes was ∼3% worse than in AFT road shoes during treadmill running at speeds between 3.9 and 5.0 m/s. Oehlert et al.^9^ showed that different AFT track spikes improved running economy by ∼2% compared to a traditional spike model during treadmill running at 14 km/h in women and at 16 km/h in men. Lab-measured changes in running economy have been shown to translate to changes in running performance,^11^ but for speeds faster than 3 m/s, percentage improvements in running speed can be expected to be smaller than observed percentage improvements in running economy.^12^ Furthermore, running economy measures are only valid when quantified during running at a sustainable, aerobic, steady-state intensity. For very fit, elite participants, this could involve running as fast as 5.8 m/s.^13,14^ However, world record pace for middle-distance events ranges from 6.2 m/s (women’s 3000 m) to 7.9 m/s (men’s 800 m). How footwear affects running economy in high-caliber runners at 5.0 m/s (e.g., ^7^) might not be indicative of how this footwear affects running performance at a speed of 7.9 m/s, almost 60% faster. Such fast middle-distance speeds require metabolic energy rates above aerobic steady-state capacity and thus quantifying physiological responses to different spikes is not feasible or valid (see ^3^).

An alternative approach to quantifying the benefits of spikes could be to have athletes perform a series of time trials in different spikes. Indeed, it has been shown previously that 3,000 m time trial performances are slower in shoes with added mass.^11^ In that study, participants ran three-time trials on an indoor track, each at the same time of day at the same day of the week, a week apart. Performing race-distance time trials at maximal effort across multiple days can profoundly interfere with athletes’ training programming and alternatively doing multiple high-effort time trials on a single day, would affect the validity of the protocol due to fatigue.

In this paper, we propose, evaluate, and use a novel approach to quantify the potential benefits of middle-distance AFT spikes. We asked athletes to perform a series of 200 m runs at self-perceived race pace (800 m or 1500 m, based on their specialty), with 10 min recovery in between, within a single session. Such 200 m race pace efforts are a common component of middle-distance training,^15,16^ and all participants were familiar with this type of workout. This is important because the premise of the proposed protocol is to measure differences in performance, while controlling effort. In a series of experiments, we compared up to four different spike models, while participants ran at least two trials in each model. First, we evaluated the validity of our novel protocol by comparing traditional spike models that were nearly identical and only differed in mass (control vs. control + 200 g) by verifying the 200 m run time differences against the theoretically expected differences. Second, we assessed the sensitivity of this novel protocol by comparing new prototype AFT spikes against traditional spikes. Third, we evaluated the reliability between three different days for two AFT spikes against traditional spikes, and finally we used this protocol to compare several commercially available AFT spikes against traditional spikes. We used an inertial measurement unit (IMU) placed at the pelvis to assess if any differences in running speed were predominantly driven by changes in step frequency or by changes in step length.

## 2. Materials and Methods

### 2.1 A Series of Experiments

This study involved four separate experiments across 3 different countries and research groups, each with 12 participants. The inclusion criteria were generally similar for all four experiments: fitting US women’s size 7, 8.5 or 9, or US men’s size 9 or 11 track spikes (Experiment 1) or fitting US men’s size 9 track spikes (Experiments 2, 3 and 4); being 16 years of age or older (Experiment 1 and 2) or 19 years of age or older (Experiment 3 and 4); currently participating in track and field training; having at minimum two years of experience training in track spikes; being free of any orthopedic, cardiovascular, or neuromuscular conditions; and having not sustained injury nor undergone any surgery within the 3 months prior to participation. Recruitment of participants was done through means of personal contacts, local track clubs, and local university track teams. Participants were Tier 3: highly-trained, national level, but not elite,^17^ with average personal bests of 2:02 in 800 to 8:55 in 3000 m across the different experiments (average World Athletics scoring tables points for each experiment: 815, 762, 817, 761, respectively (Table 1)).^18^ Because of the different timelines of the experiments and the increasing availability of AFT spikes with time, participants’ experience running with AFT varied between experiments: all participants had experience running in spikes, for Experiment 1 we only used traditional spikes, for Experiment 2 only two participants had run in AFT road shoes, none in AFT spikes, for Experiments 3 and 4 all participants had experience running with AFT road shoes and/or spikes. The characteristics of the participants in each experiment are shown in Table 1. The experiments were performed in accordance with the ethical standards of the Declaration of Helsinki. Ethics approval was obtained from the University of Massachusetts, Amherst Institutional Review Board (Protocol # 2647), the University of New Brunswick, Fredericton Research Ethics Board (REB # 2021-079), and the University of Castilla-La Mancha University Ethics Committee (Protocol CEIC924). Before taking part in the study, participants provided informed written consent.

**Table 1.** Participant characteristics for each experiment (mean ± standard deviation).

| Experiment | Participants | Age<br>(years) | Mass<br>(kg) | Height<br>(m) | World<br>Athletics<br>Score<sup>18</sup> | Location |
| --- | --- | --- | --- | --- | --- | --- |
| 1 | 8 men | 20.3 ± 2.7 | 70.9 ± 9.4 | 1.80 ± 0.06 | 787 ± 125 | US |
| | 4 women | 20.0 ± 1.4 | 56.8 ± 2.4 | 1.67 ± 0.02 | 872 ± 110 | |
| 2 | 12 men | 21.3 ± 1.8 | 62.8 ± 2.8 | 1.75 ± 0.04 | 772 ± 118 | US, Canada |
| 3 | 12 men | 27.0 ± 5.7 | 64.8 ± 3.3 | 1.76 ± 0.06 | 817 ± 126 | Spain |
| 4 | 12 men | 22.0 ± 2.7 | 69.1 ± 8.3 | 1.81 ± 0.07 | 761 ± 120 | Canada |

#### 2.1.1 Spike Conditions

For all experiments, we used the PUMA EvoSpeed Distance 9 (TRAD) as the control condition; a typical traditional track spike, lightweight in construction, while providing only minimal cushioning and no added bending stiffness elements. It has a low-profile ethylene vinyl acetate foam with a flat geometry, a midsole thickness of 17mm and 5 spike pins per shoe (Fig 1). For all spike conditions, midsole thickness was measured at the heel and the forefoot, and reported as the largest number among those locations. In Experiment 1, we compared the standard version of this spike (TRAD, 137-169 g) against the same spikes with 200 g of added mass (TRAD+, 331-368 g) (Table 2). Mass was added to the spikes by means of seven 28.3 g (1 oz) weights glued to the sides of the shoes (Fig. 1).

**Fig. 1.**
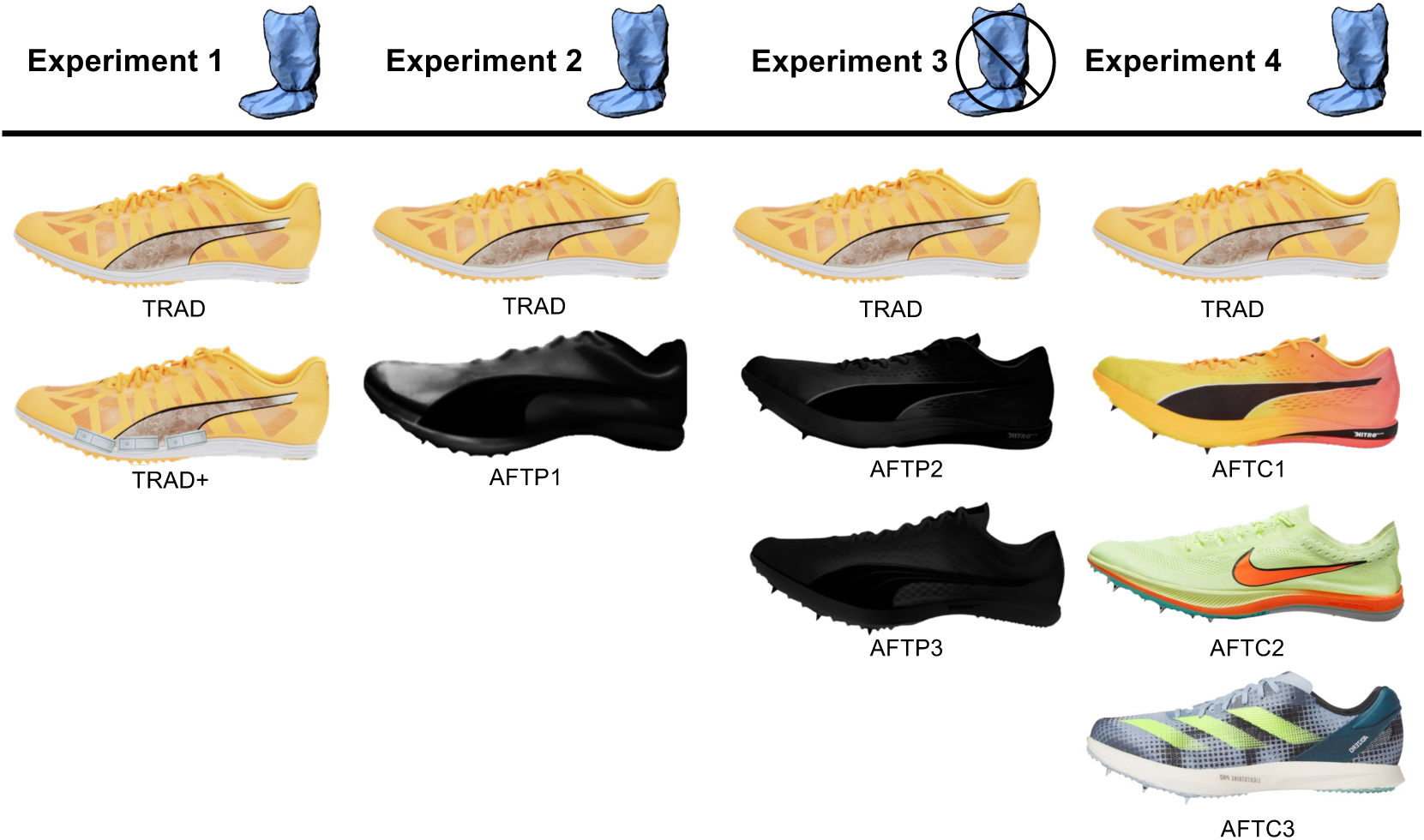
Spike shoe models used in the experiments. From top to bottom, Experiment 1: TRAD (PUMA EvoSpeed Distance 9), TRAD+ (PUMA EvoSpeed Distance 9 + 200g); Experiment 2: TRAD, AFTP1 (PUMA AFT middle-distance prototype spike); Experiment 3: TRAD, AFTP2 (PUMA AFT long-distance prototype spike), AFTP3 (PUMA AFT middle-distance prototype spike); Experiment 4: TRAD, AFTC1 (PUMA evoSPEED Long Distance Nitro Elite+), AFTC2 (Nike ZoomX Dragonfly), AFTC3 (Adidas adizero Avanti TYO).

**Table 2:** Characteristics of experimental conditions used in each Experiment.

| Spike | Sizes | Mass (g) | Midsole thickness (mm) | Experiments used |
| --- | --- | --- | --- | --- |
| TRAD | 5.5, 7, 9, 11 | 131, 149, 158, 170 | 17 | 1,2,3,4 |
| TRAD+ | 5.5, 7, 9, 11 | 331, 341, 355, 369 | 17 | 1 |
| AFTP1 | 9 | 179 | 19 | 2 |
| AFTP2 | 9 | 151 | 18 | 3 |
| AFTP3 | 9 | 149 | 19 | 3 |
| AFTC1 | 9 | 154 | 18 | 4 |
| AFTC2 | 9 | 140 | 12 | 4 |
| AFTC3 | 9 | 163 | 17 | 4 |

In Experiment 2, we compared two track spike conditions: TRAD and a PUMA AFT prototype middle-distance spike (AFTP1, 179 g). The AFTP1 spike combines a compliant, highly resilient midsole foam with an embedded carbon fiber plate. It features PUMA’s Nitro foam, a lightweight polyether block amide (PEBA) foam compound, forming a rockered midsole with a thickness of 19 mm, and 6 spike pins per shoe (Fig 1; Table 2).

In Experiment 3, we compared three track spike conditions: TRAD, a PUMA AFT prototype long-distance spike (AFTP2, 151 g) and a PUMA AFT prototype middle-distance spike (AFTP3, 149 g, different from AFTP1). The AFTP2 spike features Nitro foam, forming a rockered midsole with midsole thickness of 18 mm, but no carbon fiber plate. The AFTP3 spike combines a compliant, highly resilient midsole foam with an embedded carbon fiber plate. It features Nitro foam, forming a rockered midsole with a thickness of 19 mm, and 6 spike pins per shoe (Fig 1; Table 2).

In Experiment 4, we compared four commercially available track spike conditions: TRAD, a commercially available PUMA AFT spike (AFTC1) and two competitor brand AFT spikes (AFTC2 and AFTC3). The AFTC1 was the PUMA evoSPEED Long Distance Nitro Elite+ (154 g). It features Nitro foam, forming a rockered midsole with a thickness of 18 mm, but no carbon fiber plate, and 6 spike pins per shoe. The AFTC2 was the Nike ZoomX Dragonfly (140 g), featuring ZoomX foam, a lightweight PEBA foam compound, forming a rockered midsole with a thickness of 12 mm, but no carbon fiber plate, and 6 spike pins per shoe. The AFTC3 was the Adidas adizero Avanti TYO (163 g), featuring Lightstrike Pro foam, a lightweight thermoplastic polyether elastomer foam, forming a rockered midsole with a thickness of 17 mm, with embedded fiber glass rods for increased bending stiffness, and 6 spike pins per shoe (Fig 1; Table 2).

#### 2.1.2 Blinding Procedures

A single-blinded crossover methodology was implemented as part of the protocol for Experiments 1, 2 and 4 (i.e., participants were blinded, researchers were not). For all experiments the participants were unaware of any differences between spike conditions and which properties were modified, since the informed consent documents and researchers did not mention spike mass, foam or plate properties, only that there were several different conditions. Participants were also kept unaware of the number of different conditions or that they ran multiple times in the same condition. In none of the experiments the participants were allowed to manipulate the spikes.^11,19^ Further, for Experiments 1, 2, and 4, we used fabric boot covers (Fig. 1) such that participants did not see the spikes. These spike blinders fit around the participants spikes and lower legs in a non-restrictive fashion, providing a negligible addition to spike mass. Through an oversight, we did not use these spike blinders in Experiment 3. For this experiment, spike conditions were of a similar color/visual design, and similar to the other experiments, participants were unaware of potential differences between the spikes, and were not allowed to manipulate the spikes.

#### 2.1.3 Experiment 1 - Validation

To evaluate the external validity of our novel approach, we compared 200 m time in control spikes versus spikes that were nearly identical and only differed in mass (control vs. control + 200 g). Similar to Hoogkamer et al.,^11^ we used a mass intervention to predictably induce a change in running energetics.^20,21^ Frederick et al.^20^ showed that, across a range of speed, adding 100 g mass to each shoe increases the energy cost of running by ∼1%. The amount that a 1% increase in energy cost will slow down a runner depends on how fast they are running. Kipp et al.^12^ provided a theoretical framework supported by observations in Hoogkamer et al.^11^ that predicts that when running at 6.67m/s (200 m in 30 s), a 1% increase in energy cost will lead to a 0.6% slower run. Therefore, with 200 g added to each spike, we hypothesized that participants would run 1.2% slower.

#### 2.1.4 Experiment 2 - Sensitivity

Next, we evaluated if our protocol had the sensitivity to detect differences in what may be seen as “anecdotal” benefits of AFT spikes over traditional spikes. For this newly developed prototype middle-distance spike, we anticipated that the metabolic savings from the modern midsole foam and the embedded plate would add up to about 2%, based on preliminary running economy results from Oehlert et al.^9^ Therefore, we hypothesized that participants would run 1.2% faster in the prototype spikes.

#### 2.1.5 Experiment 3 – Reliability

We then evaluated the reliability of our novel approach to quantify the potential benefits of different AFT spikes technologies. The objective was to assess the consistency of the benefits from two AFT spikes over TRAD across three separate days. We hypothesized that the benefits of both AFT spikes would be consistent across the three measurement days.

#### 2.1.6 Experiment 4 - Commercially Available Spikes

Finally, we used our novel approach to quantify potential benefits of a set of different commercially available AFT spikes vs. traditional spikes. The aim was to evaluate if potential performance gains from AFT spikes differed between models. We hypothesized that the different AFT spikes all would result in 1.2% faster times than TRAD, with no detectable differences between AFT models. Further, we hypothesized that step frequency would be similar between conditions, while step length would be about 1% longer in the AFT spikes.^7,22^

### 2.2 Generic Experimental Protocol

#### 2.2.1 General protocol

All experiments took place at an outdoor 400 m track facility on days with little to no wind. The middle-distance track runners enrolled in the study performed a series of 200-meter runs at self-perceived 800 m or 1500 m race pace with 10 min recovery (to allow for changing spikes (seated) and walking to the start of the next trial). The effectiveness of the protocol depended on the participants’ ability to run every trial at a similar sub-maximal effort. To minimize bias, it is important that the participants are unaware of the spike intervention and that they are able to control their effort at a consistent submaximal level. Aiming to achieve the latter, we opted for a series of 200 m runs at self-perceived race pace, since this is a common component of middle-distance training,^15,16^ and all participants, like most middle-distance runners, were familiar with this type of workout, although recovery times in this experiment (10 min) were likely longer than during a real workout.

To minimize any confounding effects of participants running the first (excitement) or last (“emptying the tank”) trials at a higher effort, the number of trials that the researchers told the participants to run was greater than the number of trials they would actually run. Their first trial was considered a habituation trial and not included in the evaluation, and then after running all the needed experimental trials, the researchers revealed to the participants that the data collection was complete. Experiment 1 consisted of six experimental trials (three trials in both conditions), and we initially told participants that they would be expected to run eight, 200 m trials with 10 min recovery, but then we revealed to them that the experiment was complete after seven trials.

Similarly, for Experiments 2 and 4 we asked participants to run ten trials, but we only included four and eight experimental trials, respectively (two trials in each spike condition). For Experiment 3, the participants performed the same protocol on three separate visits, so they ran all eight trials, but we only analyzed the last six experimental trials.

The order of spike conditions was randomized for each participant prior to each session. Participants performed each 200 m trial from a standing start (similar to a middle-distance race start). Participants were unaware of the times they were running for each trial as they were not using a watch and researchers did not disclose the times they recorded until after the data collection was complete. Rest between trials was 10 minutes, to allow for changing spikes, filling out questionnaires (Experiments 2 and 4), and walking to the start of the next trial. Limited pilot data (n = 2) showed that venous blood lactate concentration did not increase over the course of an experimental session, with consistent values of ∼1.5 mmol/l before the start of the next trial.

#### 2.2.2 Experimental Setup

When the participant arrived at the track facility, we explained the study protocol to them, and they signed all the necessary forms [i.e., the Get Active Questionnaire^23^ or the Physical Activity Readiness Questionnaire (ePARmed-X+) and the informed consent document]. We asked participants to run a warmup similar to how they would for a 200 m interval workout, followed by drills and stretches as they saw fit. For Experiment 4, after their warmup, we placed an inertial measurement unit (IMU; Vicon Blue Trident sensor, Vicon, Oxford, UK) at the participant’s sacrum with a clip attached to the back of the waistband on their shorts.^24^ Next, the participant removed their shoes and researchers put fabric spike blinders on the participant’s legs (Experiments 1, 2 and 4). Then the researchers put the spikes on the participant’s feet and tightened the laces. Following this, spike blinders were pulled down over the spikes.

At the 200 m line, the researcher started the IMU recording (Experiment 4) and then moved towards the finish line. For the timing of each trial, researchers started their timers when the participant started running, indicated by them quickly lowering one hand from a raised position, and stopped their stopwatches when the participant crossed the finish line. After a trial was complete, the 200 m time was recorded and the IMU recording was stopped (Experiment 4). This was repeated for all trials.

### 2.3 Data Analyses and Statistics

We calculated average running speed in m/s by dividing the 200 m distance by the 200 m time. Raw IMU data was used to calculate spatiotemporal variables in Experiment 4. Orientation of the IMU was verified from acceleration signal by finding the gravitational component (-9.81 m/s^2^) during quiet standing before the running trial began. The vertical component was filtered using a 5 Hz cut-off frequency and multiplied by participant mass to create a vertical ground reaction force estimation, as demonstrated by Day et al.^24^ The time between peaks in the vertical ground reaction force estimate were used to calculate step time and its inverse, step frequency over the course of the full running trial. Here, step refers to the period from one foot strike to the contralateral foot strike. Average step length was calculated as the product of average speed and average step time.

We used linear-mixed effect models (LMEM) for each experiment.^25^ For all experiments, we created a LMEM with trial number as a fixed effect, participant as a random intercept, and speed as an output variable to determine if there was a significant effect of trial number on speed (i.e., slowing down or speeding up during the experiment). This was only significant for Experiment 1 (p = 0.006), so for this experiment we included trial number as a fixed effect: we created a LMEM with spike condition and trial number as fixed effects, participant as a random intercept, and speed as the output variable. For Experiment 2, we created a LMEM with spike condition as a fixed effect, participant as a random intercept, trial as a random effect, and speed as the output variable. For Experiment 3, we created a LMEM with day as a fixed effect, participant as a random intercept, and speed as an output variable to determine if there was a significant effect of day on speed. Since this was significant (p = 0.032), day was included as a fixed effect: we created a LMEM with spike condition and day as fixed effects, participant as a random intercept, trial as a random effect, and speed as the output variable. Additionally, for each day of this experiment, we created a LMEM with spike conditions as a fixed effect, participant as a random intercept, trial as a random effect, and speed was the output variable to verify that the results did not change per day. For Experiment 4, we created three LMEM with spike condition as a fixed effect, participant as a random intercept, and speed, step length, and step frequency as the output variables, respectively. A traditional level of significance (α = 0.05) was employed. Tukey HSD tests were conducted for each pairwise comparison of spike conditions. Additionally, we calculated effect size (ES) as Cohen’s d and reported percent differences for each measure with the intention of providing relevant context for our determined p values. ES with values of 0.2, 0.5, and above 0.8 were considered as small, medium, and large, respectively.^26^ To evaluate the reliability, in Experiment 3 we calculated the coefficient of variation (CV = SD / mean × 100%) within each day and for each spike, across days. The ranges of values considered for categorizing CV% were as follows: CV < 10 as very good, 10–20 as good, 20–30 as acceptable, and CV > 30 as unacceptable.^26^ Further, for each participant we calculated the average speeds in each spike across days 1 and 2, days 1 and 3, days 2 and 3, and across days 1, 2 and 3. Then, we calculated the speed difference between the TRAD and the AFTP3 spikes. Next, we calculated Pearson’s correlation coefficients, for these differences, between each individual day and the 3-day average, and between each 2-day average and the 3-day average. Correlation coefficients with values of 0.1, 0.3, and 0.5 were considered as small, medium, and large, respectively, with 1.0 being a perfect correlation^26^. Statistical analyses were performed in R using lme4 package^27^ for the linear mixed effect models and emmeans package^28^ for post hoc Tukey HSD tests. Descriptive statistics are reported as mean ± standard deviation (SD). For visual display in the figures, we averaged the 200 m trial times that were performed in the same spike condition for each experimental session (three trials for Experiment 1; two trials for Experiments 2, 3 and 4).

## 3 Results

In Experiment 1, participants ran significantly slower in the TRAD+ spikes (5.97 ± 0.52 m/s) than in the TRAD spikes (6.04 ± 0.50 m/s) (1.2%, ES = 0.14, p = 0.016, n = 12; Fig. 2). Participants significantly slowed down during the experimental session (0.05 m/s per set of two trials, p = 0.004). In Experiment 2, participants ran significantly faster in the AFTP1 spikes (6.35 ± 0.42 m/s) than in the TRAD spikes (6.28 ± 0.50 m/s) (1.1%, ES = 0.15, p = 0.030, n = 12; Fig. 2).

**Fig. 2.**
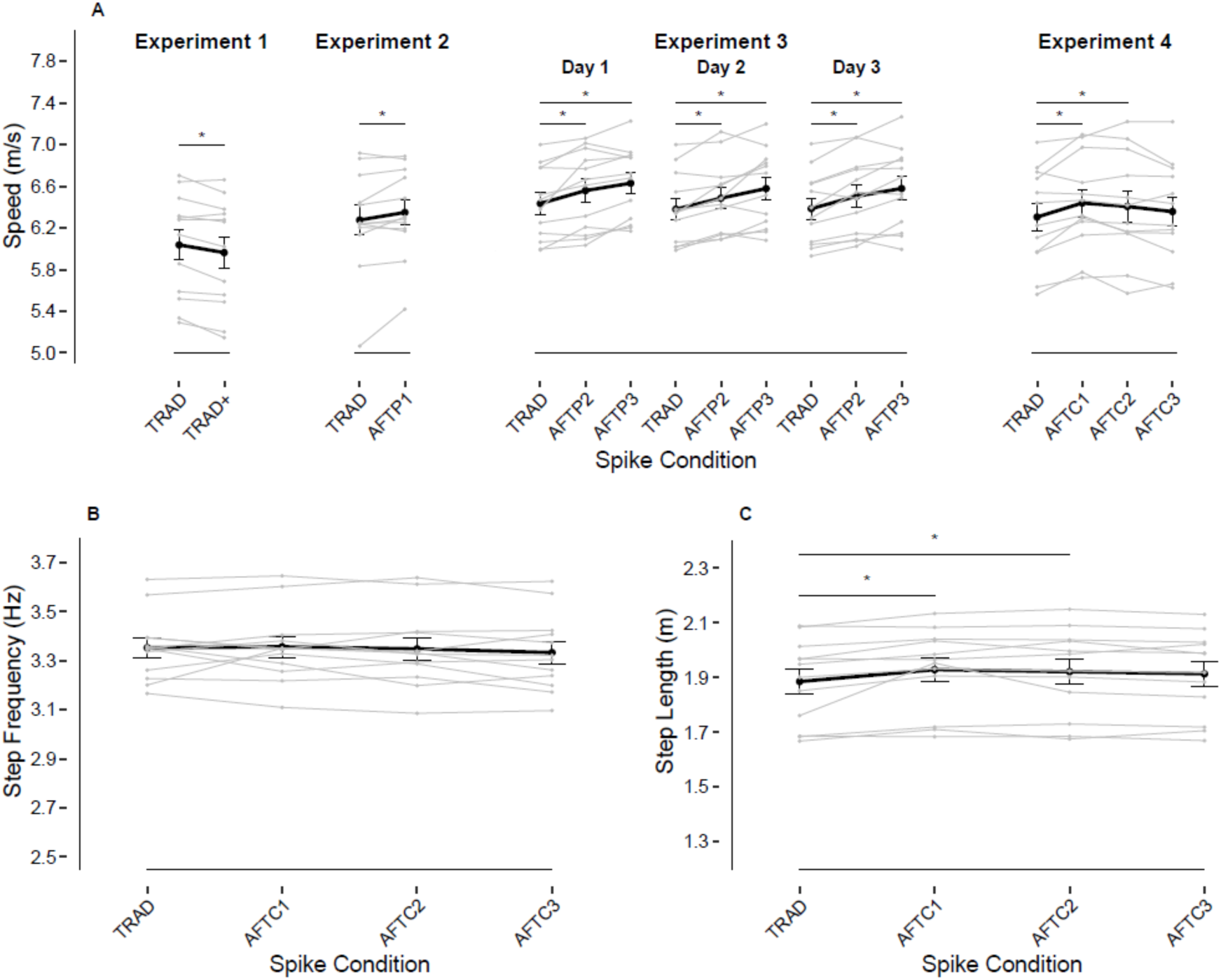
Spike conditions significantly affect running speed and step length, but not step frequency. Panel **A** shows average running speeds in different spike conditions, for Experiment 1, 2, 3 and 4, from left to right. Panel **B** shows step frequency was similar across spike conditions. Panel **C** shows significantly longer step lengths for

In Experiment 3, participants ran significantly faster in the AFTP2 spikes (6.52 ± 0.37 m/s) and the AFTP3 spikes (6.60 ± 0.36 m/s) than in the TRAD spikes (6.40 ± 0.34 m/s) (1.8%, ES=0.34, p < 0.0001 and 3.1%, ES= 0.57, p < 0.0001, respectively, n = 12; Fig. 2). Running speeds in AFTP3 spikes were significantly faster than in AFTP2 spikes (1.2%, ES=0.19, p = 0.0003). The between-spike results were consistent between days. The CV% values were classified as very good for inter-session (5.5%, 5.4% and 5.7% for day 1, 2 and 3 respectively) and intra-session (5.3%, 5.5% and 5.5% for TRAD, AFTP2 and AFTP3 respectively). The linear mixed effects model analyses for each day separately all showed similar differences between spikes, both in magnitude (% change and effect size) and in significance (Table 3).

**Table 3:** Magnitude and significance of between spike differences were.

| Comparison |  | Day 1 | Day 2 | Day 3 | 3-Day average |
| --- | --- | --- | --- | --- | --- |
| TRAD vs. AFTP2 | % change | 1.9 | 1.7 | 1.9 | 1.8 |
|  | effect size | 0.36 | 0.32 | 0.31 | 0.34 |
|  | p value | 0.003 | 0.004 | 0.0003 | <0.0001 |
| TRAD vs. AFTP3 | % change | 3.0 | 3.1 | 3.0 | 3.1 |
|  | effect size | 0.57 | 0.56 | 0.51 | 0.57 |
|  | p value | <0.0001 | <0.0001 | <0.0001 | <0.0001 |
| AFTP2 vs. AFTP3 | % change | 1.1 | 1.4 | 1.1 | 1.2 |
|  | effect size | 0.19 | 0.25 | 0.21 | 0.19 |
|  | p value | 0.124 | 0.015 | 0.035 | 0.0003 |

The speed difference between TRAD and AFTP3 for each individual participant varied across days. Pearson’s correlation coefficients between day 1, day 2 and day 3, and the 3-day average were large: 0.80, 0.77, and 0.89, respectively. However, the Pearson’s correlation coefficients between each 2-day average and the 3-day average were substantially higher: 0.97, 0.90 and 0.96 for day 1 and 2, day 1 and 3, and day 2 and 3, respectively.

In Experiment 4, participants ran significantly faster in the AFTC1 spikes (6.44 ± 0.45 m/s) and the AFTC2 spikes (6.41 ± 0.51 m/s) than in the TRAD spikes (6.31 ± 0.47 m/s) (2.1%, ES= 0.31, p = 0.002 and 1.6%, ES= 0.22, p = 0.035, respectively, n = 12; Fig. 2). Running speeds in the AFTC3 spikes (6.36 ± 0.48 m/s) were not significantly different from the TRAD spikes (0.8%, ES=0.13, p = 0.493). Running speeds were not significantly different between the AFTC spikes (AFTC1 vs. AFTC2 p = 0.800; AFTC1 vs. AFTC3 p = 0.116; AFTC2 vs. AFTC3 p = 0.534), but the numerical difference between AFTC1 and AFTC3 was more than 1% (1.3%, ES = 0.17).

Step frequency was not different between spikes (all p > 0.52; Fig. 2), but participants ran with significantly longer steps in AFTC1 spikes (1.93 ± 0.15 m) and AFTC2 spikes (1.92 ± 0.16 m) than in the TRAD spikes (1.88 ± 0.15 m) (2.3%, ES = 0.33, p = 0.001; 1.9%, ES = 0.26, p = 0.008; Fig. 2). Step lengths in the AFTC3 spikes (1.91 ± 0.15 m) were not significantly different from the TRAD spikes (1.5%, ES = 0.20, p = 0.0512). Step lengths were not significantly different between the AFTC spikes (AFTC1 vs. AFTC2 p = 0.893; AFTC1 vs. AFTC3 p = 0.472; AFTC2 vs. AFTC3 p = 0.875).

## 4 Discussion

In this paper, we proposed, evaluated and used a novel approach to quantify the potential benefits of middle-distance AFT spikes. The combined results indicate that our novel approach has the validity, sensitivity and reliability to quantify the benefits of middle-distance AFT spikes. The results of Experiment 1 provide insights into the validity of our interval-based approach. On average, participants ran 1.2% slower in the spikes with 200 g added mass, similar to the 1.2% difference we hypothesized. The results of Experiment 2 indicate that our approach has the sensitivity to confirm what so far has been considered “anecdotal” benefits of AFT spikes over traditional spikes.

Participants ran significantly (1.1%) faster in AFT prototype spikes than in traditional spikes. The results of Experiment 3 indicate that our approach is reliable. Across three separate days, the between-spike differences in running speed were similar, both in magnitude (% change and effect size) and in significance. However, at the level of individual runners, the responses to AFT spikes varied between days. Our data suggests that two separate sessions are needed to obtain reliable individual data. Finally, we used our interval-based approach to compare several commercially available AFT spikes against traditional spikes. In two AFT spike models participants ran significantly (2.1% and 1.6%) faster than in the traditional spikes. Speed was similar between a third AFT spike model and the traditional spikes. These speed results were mirrored by changes in step length, as participants took significantly (2.3% and 1.9%) longer steps in the two faster AFT spike models, while step length was similar between the third AFT spike model and the traditional spikes.

### 4.1 Construct Validity

Shoe mass is one of the most studied features of running shoes.^11,20,21,29^ There is currently a strong consensus that there is a linear relationship between the increase in shoe mass and the increase in running energy cost, with an increment in the oxygen/energy cost of ∼1% per added 100 g of shoe mass.^20^ Kipp et al.^12^ provided a theoretical framework to translate changes in energy cost into changes in running performance, based on the known curvilinear relationship between running speed and energy cost and the cost of overcoming air resistance, and supported by shoe-mass time trial results from Hoogkamer et al.^11^ Using this framework, we anticipated that participants would run 1.2% slower in the TRAD+ spikes with 200 g added mass. We found that on average our participants ran 1.2% slower, with 75% of the participants (9 of 12) running slower in the TRAD+ spikes. Our participants ran slightly slower than anticipated (∼6.0 m/s vs. 6.67 m/s) and at that speed the framework predicts a 1.3% slower speed in the TRAD+ spikes (not 1.2% slower). While our results are close to our prediction, it should be acknowledged that our predictions are based on the 1% for 100 g shoe mass rule of thumb and that this 1% number slightly varies between studies,^11,20,21,29^ and that it might be less than 1% at faster speeds.^20^

### 4.2 Sensitivity

Next, we evaluated if our interval-based approach is sensitive enough to detect differences between AFT spikes and traditional spikes. We anticipated that speed differences between AFT spikes and traditional spikes would be similar to the effect of 200 g added mass and we hypothesized that participants would run 1.2% faster in the

AFT prototype spikes. Participants ran 1.1% faster in the AFT prototype spikes than in the traditional spikes, which is equivalent to the effect of reducing the mass of the traditional spikes by ∼180 g per spike (which is not possible, considering that the traditional spikes weigh ∼150 g each).

### 4.3 Reliability

We evaluated the reliability of this protocol by asking 12 participants to run in the same three spike conditions with the same protocol on three separate days. We found that, at the group level, results were similar between all three days (Table 3) and similar to the 3-day average values (significant speed improvements of 1.8 and 3.1% of the AFTP2 and AFTP3, compared to TRAD, respectively). The Pearson correlation coefficients for Day 1, 2, and 3 vs. the 3-day average were large (0.77-0.89), but the correlation coefficients for 2-day average vs. 3-day average results were substantially higher (0.90-0.97), suggesting that a minimum of two days is needed to quantify dependable individual responses to AFT spikes. Quantifying individual responses to AFT shoes is common when aiming to identify biomechanical mechanisms underlying improvements from AFT,^30,31^ or characteristics of runners who experience more or less benefits.^32–34^ However, as discussed by Barrons et al.,^35^ such analyses critically depend on the accuracy of the response measurement. Note that both for running economy measures^35^ and for speed outcomes from our interval-based approach, multiple sessions with repeated conditions are recommended, when attempting to quantify individual responses to AFT. In our study, 75% of the participants ran consistently faster in AFTP2 vs. TRAD across the three days, and similarly 75% of the participants ran consistently faster in AFTP3 vs. TRAD across the three days (Fig. 3). For AFTP2 vs. TRAD, two participants (17%) ran faster in two of the three days, and one participant (8%) only ran faster on one day. For AFTP3 vs. TRAD, three participants (25%) ran faster on two of the three days. Nine participants (75%) ran faster in AFTP3 vs. AFTP2, and eight of them did this consistently across the three days (day 1 for participant #6 being the exception). For the three participants who ran faster in AFTP2 vs. AFTP3 (#8, 9 and 10), two of them did this consistently across the three days (day 1 for participant #8 being the exception). Noteworthy, the footwear condition with plate (AFTP3) lead to significant improvements over the condition with similar construction without plate (AFTP2). This is in contrast to Healey and Hoogkamer^36^ who observed that making medio-lateral cuts in the carbon- fiber plates of AFT shoes (i.e., effectively reducing the shoes’ longitudinal bending stiffness) did not significantly worsen running economy. This discrepancy might be related to the different speeds run in each experiment (∼6.50 m/s for Experiment 3 vs. 3.89 m/s in Healey and Hoogkamer ^36^), suggesting that increased longitudinal bending stiffness is more beneficial at faster running speeds.

**Fig. 3.**
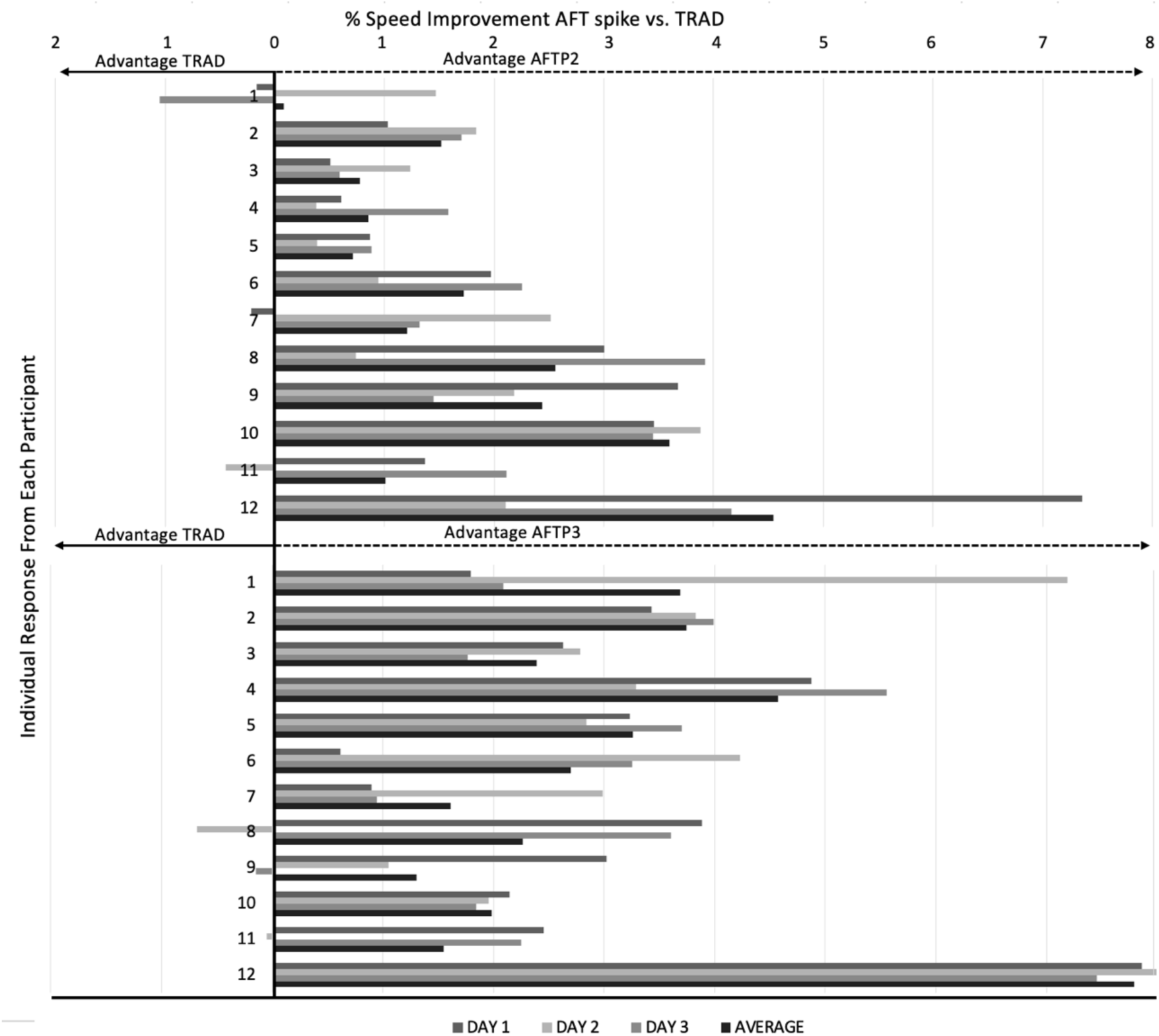
Individual response of each participant in prototype AFT spikes vs. traditional spikes (TRAD) in Experiment 3 for each of the three days and the 3-day average.

### 4.4 Quantifying the Benefits of AFT Spikes

Finally, we used our novel approach to quantify potential benefits of several different commercially available AFT spikes over traditional spikes. Participants ran significantly faster in the AFTC1 (2.1%) and AFTC2 (1.6%) than in the TRAD spikes. However, running speeds were not significantly different in the AFTC3 spikes versus the TRAD spikes (0.8%), nor between the AFTC spikes. The improvements from the commercially available AFT spikes compared to the traditional spikes were similar to, but slightly less than, those of the prototype AFT spikes evaluated in Experiment 3 of this study (1.8 - 3.1%). Over 800 or 1500 m at these speeds, athletes would run 2.5 or 5 sec faster, respectively, in these commercially available AFT spikes than in traditional spikes.

While the effects of spikes on sprinting performance have been studied for at least 50 years, with an inquiry into the potential benefits of the banned Puma brush spikes in the early seventies^37^, here we specifically focused on middle-distance running spikes. Given the novelty of AFT spikes, and the challenges of quantifying their potential benefits on middle-distance running performance,^3^ we are not able to directly compare our findings to any previous studies in the literature.

Willwacher et al.^8^ performed a longitudinal, observational, and retrospective study design evaluating the 100 best performances per year for men and women in outdoor track events from 2010 to 2022. They observed more pronounced improvements for the 1500 m and longer distances (0.7 - 1.8%) than shorter distances (100 m to 800 m: 0.3-0.6%) for women, and less of an effect of AFT spikes in men. The differences at the group level in the 800 m were ∼0.7 sec for both men and women, and ∼1.3 and ∼2.2 sec in the 1500 for men and women, respectively. Substantially smaller than the 2.5 and 5 sec time savings that our data indicate. When comparing running economy results for AFT spikes at 4.4 and 5.0 m/s, Oehlert et al.^9^ found 1.8 and 2.1% improvements for two AFT spike models as compared to traditional spikes. Such running economy improvements can be expected to improve middle-distance running performance by ∼1.2%,^12^ which is less than the speed improvements we observed in the current study at ∼6.4 m/s. Although, not specific to AFT spikes, some earlier studies have evaluated 3,000 m time trial performance on the track in AFT road shoes and reported improvements ranging from 1 to 5%, similar to our spike results.^33,38^ Without mechanical characterization of midsole compliance and resilience, and longitudinal bending stiffness of the different commercial AFT spike models (beyond the scope of this study), we can only speculate why participants only ran significantly faster in AFTC1 and AFTC2 vs. TRAD, and not in AFTC3. It might be that the compliance and resilience properties of the thermoplastic polyether elastomer foam in the AFTC3 are less than that of the PEBA-based foams in AFTC1 and AFTC2, or that the bending stiffness of AFTC3 is suboptimal. Importantly, the exact benefits of increased longitudinal bending stiffness within AFT shoes are still under debate,^39,40^ with some evidence that the improved foam properties are more important.^22,36,41^

Across Experiment 2, 3 and 4, participants consistently ran faster in the AFT spikes than in the traditional spikes (with the exception of AFTC3). The three experiments each used slightly different models of AFT spikes and the group-level benefits varied from 1.1% in Experiment 2, 1.8 and 3.1% in Experiment 3 to 2.1 and 1.6% in Experiment 4. The consistency of these findings, while these experiments were performed by different research teams in three different countries (US, Canada and Spain) is a testament of the robustness of our interval-based approach and further supports the validity of the observed benefits of AFT spikes.

Intriguingly, the faster speeds in the AFTC1 (2.1%) and AFTC2 (1.6%) spikes as compared to the TRAD spikes in Experiment 4, were accompanied by 2.3% (i.e., 4.3 cm) and 1.9% (i.e., 3.6 cm) longer steps in the two faster AFTC1 and AFTC2 spikes, respectively. Step lengths were similar between the other spike models. Furthermore, step frequency was similar across all spikes (∼3.35 Hz, or 201 steps per minute). Combined, this suggests that the performance benefits from AFT spikes in middle-distance running originate from the spikes allowing runners to take longer step lengths without requiring any compensation in step frequency. This is supported by observations of runners taking longer steps at fixed speeds in AFT road shoes vs. traditional shoes.^7,22,42^ To confirm this, future studies could compare step lengths in AFT spikes vs. traditional spikes for running at a fixed, middle-distance running speed. With average step lengths of 1.93 m and 1.92 m in the AFTC1 and AFTC2 spikes, the participants took only 103.6 steps and 104.1 steps during their 200 m, respectively, about 2.5 steps less than in the traditional spikes. Over a 1500 m race this would add up to 21 or 17 less steps in the AFTC1 and AFTC2 spikes than in traditional spikes, respectively.

These biomechanical changes are similar to those reported in treadmill studies on AFT road shoes but different from preliminary findings for AFT spikes. Treadmill studies of road shoes consistently report that runners take ∼1% longer steps in AFT shoes than in traditional shoes.^7,10,22,42^ Since running speed is fixed between conditions in such studies, these longer steps are accompanied by reductions in step frequency. In a case study on overground, track running, Russo et al.^43^ evaluated biomechanics during 80 m trials at 95% of maximal speed comparing AFT spikes with a carbon-fiber plate and a modern midsole foam against traditional spikes. The athlete ran at the same speed (∼8.0 m/s) and did not substantially change step length or step frequency, even though contact time was longer in the AFT spikes. The fixed speed and single subject nature of Russo et al.^43^ might explain the differences with our study.

### 4.5 Limitations and Future Directions

While our approach of evaluating a series of 200 m runs at a self-perceived race pace, interferes less with athletes’ training programming and is less sensitive to day-to-day variability than performing race-distance time trials at maximal effort across multiple days, self-perceived sub-maximal efforts are not a direct measure of running performance and sensitive to bias. For Experiments 2-4, it may be possible that the participants ran faster in the AFT spikes for 200 m simply because the AFT spikes felt bouncier or more comfortable, or because of their belief in performance enhancement from AFT spikes.

These faster sub-maximal 200 m times might not translate into any true performance gains over a full 800 m or 1500 m race at maximal effort. However, results for Experiment 1 show that participants quite precisely adjust their speed according to the expected metabolic energy increase from the added mass, despite the identical underfoot construction of the two spike conditions (TRAD and TRAD+). Furthermore, our participant blinding procedures were effective in that participants were typically unaware of how many different spike conditions they were running in, and what spikes they were currently wearing. In general, any speculation by the participants about their shoe conditions were inaccurate, and researchers neither confirmed nor denied their comments. Future further proof of validity could come from experiments that directly compare between-spike differences observed with our interval-based approach to between-spike differences in race distance time trials.

For researchers looking to implement our interval-based protocol in future studies, great care must be taken to limit the bias of the researchers and participants. Participants should be blinded to the shoe conditions, not just to what is currently on their feet. If, for example, the participant knows that they will be wearing one traditional spike and one AFT spike, the participant might be able to use this information to correctly deduce the spike condition they are currently wearing. With AFT spikes becoming more common, especially when comparing commercially available spikes, some participants might be able to feel or recognize differences in midsole cushioning or longitudinal bending stiffness between spikes, which could bias their performances in our effort-based protocol. We recommend that investigators mirror the order of spike conditions (e.g., ABCCBA) and counterbalance condition orders across participants (i.e., each possible order is done, and all orders are done by a same number of participants), to minimize any order effects. Participants ran their 200 m trials at consistent speeds throughout the test session for Experiments 2-4, but in Experiment 1 they slowed down significantly from the first set of trials in each condition to the third set of trials in each condition.

While our study shows that our novel interval-based approach to quantify potential benefits of middle-distance AFT spikes is valid and reliable at the group level, we caution against using the results of a single session to label an individual as a responder/non-responder to AFT spikes, or to compare differences in responses between individuals. Specific to that purpose, we found that at least two days are necessary to be able to evaluate individual responses to AFT spikes.

Additionally, there are limitations with the implementation of this protocol. In this current study, running times were recorded by hand, which is subject to human error.

Step lengths were calculated as the average of the 200 m run, based on the average step frequency and the running time. Future studies could place IMUs at the feet, which would allow accurate calculation of stride length at a stride-by-stride resolution.^44^ Evaluating between-spike differences in stride length at a stride-by-stride resolution could provide additional information about how benefits of AFT spikes potentially vary from the acceleration phase to the top speed phase.

Finally, caution should be taken when interpreting the results on the benefits of AFT spikes in Experiment 4. On average, the male participants in this experiment ran their trials at ∼6.4 m/s, which is equivalent to a 2:05 min 800 m, or 3:54 min 1500 m. To evaluate the benefits at elite or even world record level, this experiment should be repeated with elite runners. Still, our findings at ∼6.4 m/s are likely more informative than running economy measurements for running at 16 km/h (4.44 m/s; equivalent to a 3:00 min 800 m). Elite and/or female runners might have different plantar flexor strength or body mass than our male participants, factors that are likely to affect responses to AFT.^45,46^ At the start of our study, we only had spikes available in the men’s sample size US9 (except for Experiment 1), but follow-up studies should include more female participants.

## 5 Conclusions

To overcome the challenges in quantifying the potential benefits of advanced footwear technology spikes related to the physiology of middle-distance running intensity, we introduced a novel interval-based approach consisting of a series of 200 m runs at self-perceived middle-distance race pace. In a series of experiments, we showed the validity and reliability of our method and then used it to compare commercially available spikes. Our reliability experiment showed that group-level results were consistent across days, but that at least two separate sessions are needed to evaluate individual responses. Participants ran ∼2% faster and with ∼2% longer steps in two commercially available AFT spike models than in traditional spikes, while speed and step length were similar between a third AFT spike model and the traditional spikes.

## Acknowledgements

We thank the athletes of Athletics Canada and the UNB Reds Varsity Track and Cross Country team, and of the running communities in Amherst, Boston, San Antonio, Castilla La-Mancha and Madrid for their participation in our study. We thank Rodger Kram and Jose van der Veen for energetic(s) discussions to spark the ideas presented in this paper. We thank Dan Feeney for guidance regarding our statistical analysis. Lastly, we thank our undergraduate researchers, Elizabeth Hoenes, Caleb Zylka, Violeta Muñoz de la Cruz and Robert Carew for assisting with data collections and recruitment. All spikes were provided by PUMA SE.

## Declarations

**Compliance with Ethical Standards**

### Ethical approval

The study was performed in accordance with the ethical standards of the Declaration of Helsinki. Ethics approval was obtained from the University of Massachusetts Institutional Review Board (Protocol# 2647), from the University of New Brunswick Research Ethics Board (REB #2021-079) and the University of Castilla-La Mancha University Ethics Committee (Protocol CEIC924)

### Informed consent

Written informed consent was obtained from all individual participants included in the study.

## Funding

This research was partly supported by a research contract from PUMA SE with the University of Massachusetts, Amherst. All spikes were provided by PUMA SE.

## Conflicts of interest / Competing interests

Montgomery Bertschy, Victor Rodrigo-Carranza, Ethan W.C. Wilkie, Jeremy Noble and Wayne J. Albert have no conflicts of interest relevant to the content of this article.

Wouter Hoogkamer is a paid consultant to PUMA and has received research grants from PUMA and Saucony. Laura A. Healey is an employee of PUMA. No footwear company had any influence on the conceptualization of this study or results presented in this publication.

## Data availability

All data discussed in this manuscript are provided in the tables.

## Authors’ contributions

Conceptualization: Wouter Hoogkamer; Experimental design: Montgomery Bertschy, Laura A. Healey, Wouter Hoogkamer; Data collection: Montgomery Bertschy, Victor Rodrigo-Carranza, Ethan W.C. Wilkie, Jeremy Noble; Data analysis: Montgomery Bertschy, Victor Rodrigo-Carranza, Ethan W.C. Wilkie, Jeremy Noble, Wayne J. Albert; Writing - original draft preparation: Montgomery Bertschy, Victor Rodrigo-Carranza, Ethan W.C. Wilkie, Laura A. Healey, Wouter Hoogkamer; Writing - review and editing: Montgomery Bertschy, Victor Rodrigo-Carranza, Ethan W.C. Wilkie, Laura A. Healey, Jeremy Noble, Wayne J. Albert, Wouter Hoogkamer

## Notes

### Summary of Updates

Minor wording changes throughout; Discussion updated; Description of Figure 2 revised.

